# Methionine oxidation of TRPV2 regulates thermogenesis in brown adipocytes

**DOI:** 10.1101/2025.05.02.651863

**Authors:** Mari Iwase, Satoko Kawarasaki, Tsuyoshi Goto, Kunitoshi Uchida

**Author notes:** To whom correspondence should be addressed: Mari Iwase, Ph.D, and Kunitoshi Uchida, Ph.D Laboratory of Functional Physiology, Department of Environmental and Life Sciences, School of Food and Nutritional Sciences, University of Shizuoka, Yada 52-1, Suruga-ku, Shizuoka, Shizuoka 422-8526, JAPAN, (M.I) and (K.U.).

## Abstract

Transient receptor potential vanilloid 2 (TRPV2) is a non-selective cation channel activated by mechanical stimuli and temperatures above 52°C. Although we have previously reported that TRPV2 regulates non-shivering thermogenesis through facilitating the expression of genes related to thermogenesis, how TRPV2 activity is regulated in brown adipocytes under physiological conditions remains unclear. Recently, it was reported that methionine oxidation was shown to reduce the temperature threshold of TRPV2 for activation to core body temperature or lower. In the present study, we investigated whether methionine oxidation activates TRPV2 and regulates thermogenesis in the differentiated brown adipocytes. As a result, treatment with Chloramine-T (ChT, a methionine oxidant) activated TRPV2 at temperatures of >30°C in mouseTRPV2-expressing HEK293T cells and in the differentiated brown adipocytes. Moreover, ChT treatment enhanced the expression of genes related to thermogenesis in the differentiated brown adipocytes. These results suggest that methionine oxidation might activate TRPV2 at around body temperature and increase thermogenesis-related gene expression.

## INTRODUCTION

Adipose tissues are classified as white adipose tissues (WAT) and brown adipose tissues (BAT) according to their functions. WAT stores the energy as triglycerides and BAT expends the energy as heat. This thermogenesis in BAT liberates heat via mitochondrial uncoupling protein 1 (UCP1) in the inner mitochondrial membrane (Cannon and Nedergaard, 2004). UCP1 expression is induced by the sympathetic nervous system (SNS) signaling, which is activated by cold exposure and intake of specific food ingredients. Noradrenaline released from sympathetic nerve endings binds to β3-adrenergic receptors and activates intracellular adenylate cyclase, increasing intracellular cyclic AMP (cAMP) levels. The increase in cAMP activates protein kinase A (PKA), upregulating *Ucp1* expression via activation of cAMP response-element binding protein, peroxisome proliferator-activated receptor γ (PPARγ) coactivator α (PGC1α), and activating transcription factor 2 (Tabuchi and Sul, 2021). PKA activation also facilitates the lipolysis of lipid droplets to provide free fatty acids to produce heat (Fredriksson et al., 2001). Although thermogenesis in BAT is essential for maintaining body temperature in infants, its function decreases with age. This is considered one of the causes of age-related obesity, and elucidation of the thermogenesis mechanism in BAT is expected to prevent and improve obesity and related diseases (Lidell et al., 2014).

Redox states in the body are maintained by intracellular antioxidants and redox substances such as reactive oxygen species (ROS). Disruption of this homeostasis causes damage to DNA, proteins, and lipids that induce cancer growth and neurodegenerative diseases such as Alzheimer’s disease and Parkinson’s disease (Didier et al., 2023). In BAT, ROS accumulation causes oxidative stress, leading to mitochondrial dysfunction and decreased thermogenesis (Shadel and Horvath, 2015). Hydrogen peroxide (H_2_O_2_) treatment of brown adipocytes impaired their thermogenesis, alleviated by co-treatment with antioxidants such as vitamin E (Cui et al., 2019). These reports indicated that oxidative stress negatively regulates BAT thermogenesis. In contrast, ROS may activate thermogenesis by altering thermogenesis-related gene expression or by the direct redox modification of UCP1. It was reported that ROS promoted UCP1 expression by amplifying cAMP-induced p38 mitogen-activated protein kinase activation in BAT after cold stimulation in mice (Ro et al., 2014). Cold stimulation increased mitochondrial ROS levels, contributing to the sulfenylation of cysteine 253 in UCP1 and activation of UCP1-mediated thermogenesis in brown adipocytes (Chouchani et al., 2016; Shi et al., 2021; Oo et al., 2022). Although ROS have been shown to modulate thermogenesis through various mechanisms, many aspects of their actions remain unclear, and the role of methionine oxidation in brown adipocytes is even not well understood.

Transient receptor potential vanilloid 2 (TRPV2) is a Ca^2+^-permeable non-selective cation channel activated by noxious heat with an activation temperature threshold higher than 52°C (Caterina et al., 1999). Some chemical ligands such as 2-aminoethoxydiphenyl borate (2APB) and lysophosphatidylcholine also activate TRPV2 in a species-specific manner (Juvin et al., 2007; Monet et al., 2009). TRPV2 has been reported to detect cell stretching and swelling via its mechanosensitive properties (Muraki et al., 2003; Shibasaki, 2016). TRPV2, which is expressed in the central and peripheral nervous systems, macrophages, pancreas, and cardiovascular systems, plays important roles in axon outgrowth in developing neurons (Shibasaki et al., 2010), intestinal movement (Mihara et al., 2010), phagocytosis (Nagasawa et al., 2007), insulin secretion (Sawatani et al., 2019), and maintenance of cardiac structure and function (Muraki et al., 2007). We previously reported that TRPV2 is expressed in brown adipocytes and that its expression increases during differentiation (Sun et al., 2016b). The activation of TRPV2 also impaired the differentiation of brown adipocytes (Sun et al., 2016b). In addition, TRPV2 positively regulates non-shivering thermogenesis in differentiated brown adipocytes (Sun et al., 2016a). Since the core body temperature is not generally exposed above 52°C, it has not been well clarified how the activity of TRPV2 in brown adipocytes is modulated under physiological conditions. Interestingly, Fricke et al. reported that rat and human TRPV2 were activated by treatment with a methionine oxidant at 32°C (Fricke et al., 2019), indicating that oxidized TRPV2 could be activated by the physiological range of temperature. However, the physiological significance of TRPV2 oxidation has not yet been elucidated.

In the present study, we hypothesized that methionine oxidation of TRPV2 might be involved in thermogenesis. We showed that treatment with a methionine oxidant enables the activation of TRPV2 by the physiological range of temperature and enhances the expression of the genes related to thermogenesis in brown adipocytes.

## MATERIALS & METHODS

### HEK293T cells

Human embryonic kidney-derived 293T (HEK293T) cells were maintained in DMEM (FUJIFILM Wako Pure Chemical Corp., Osaka, Japan) containing 10 % FBS (Thermo Fisher Scientific Inc, Massachusetts, USA), 50 units/mL penicillin (Thermo Fisher Scientific Inc), 50 μg/mL streptomycin (Thermo Fisher Scientific Inc), and 2 mM L-glutamine (GlutaMAX, Thermo Fisher Scientific Inc) at 37°C in 5 % CO_2_. For Ca^2+^-imaging, 1 μg of plasmid DNA containing mouse TRPV2 (mTRPV2) in pcDNA3.1 in FBS- and antibiotics-free DMEM medium (FUJIFILM Wako Pure Chemical Corp) were transfected to HEK293T cells using Lipofectamine Plus Reagent (Thermo Fisher Scientific Inc). After 3 to 4 hours incubation, cells were reseeded on coverslips and further incubated at 37°C in 5 % CO_2_. Ca^2+^-imaging was performed one day after transfection.

### Brown adipocytes line

We used immortalized pre-adipocytes isolated from interscapular BAT of UCP1-mRFP1 transgenic mice (Kawarasaki et al., 2019; Kenmochi et al., 2022). These cells were cultured in standard medium containing 10 % FBS, 100 units/mL penicillin, and 100 μg/mL streptomycin in DMEM at 37°C in a 5 % CO_2_. After reaching confluence, immortalized pre-adipocytes were differentiated with standard medium supplemented with 10 μg/mL insulin (FUJIFILM Wako Pure Chemical Corp.), 1 nM triiodothyronine (T_3,_ Thermo Fisher Scientific Inc.), 0.125 mM indomethacin (FUJIFILM Wako Pure Chemical Corp.), 0.25 μM dexamethasone (Thermo Fisher Scientific Inc.), and 0.5 mM 3-isobutyl 1-methylxanthine (Nacalai tesque Inc. Kyoto, Japan) for 48 h. The media were replaced every 2 days with differentiation medium consisting of standard medium supplemented with 5 μg/mL insulin and 1 nM T_3_. Ca^2+^-imaging, reverse-transcription polymerase chain reaction (RT-PCR) and RT-qPCR were performed after 8 days differentiation.

### Ca^2+^-imaging

HEK293T cells on coverslips were mounted in an open chamber. Immortalized pre-adipocytes were differentiated on glass bottom dish (Matsunami Glass Ind., Ltd.). This open chamber with coverslips and glass bottom dish were superfused with a standard bath solution (140 mM NaCl, 5 mM KCl, 2 mM MgCl_2_, 2 mM CaCl_2_, 10 mM HEPES, and 10 mM glucose, pH 7.4). All chemicals were dissolved in the standard bath solution. Standard bath solution was perfused using peristaltic pump at a rate of 3.5 ml/min (ELEYA SMP-23; Tokyo Rikakikai Co., LTD., Tokyo, JAPAN). Thermal stimulation was applied by increasing the bath temperature with an inline heater (SH-27B; Warner Instruments, Massachusetts, USA). Intracellular Ca^2+^ concentrations ([Ca^2+^]_i_) in these cells were measured by dual-wavelength Fura-2 microfluorometry (excitation at 340/380 nm and emission at 510 nm, Thermo Fisher Scientific Inc) and the CoolSNAP ES CCD camera (Photometrics, Arizona, USA) and recorded using NIS Elements v4.50 software (NIKON Corp., Tokyo, Japan) at 5-s intervals. The ratio image was calculated using ImageJ (NIH).

### Reverse-transcription polymerase chain reaction (RT-PCR)

The differentiated brown adipocytes were subjected to 3 μM Isoprenaline and/or 1 mM Chloramine T (ChT), and 3 μM Isoprenaline (Iso) and/or 3 mM Dithiothreitol (DTT) for 4 h. Total RNA was purified from the brown adipocytes using NucleoSpin RNA (Macherey–Nagel GmbH & Co., Duren, Germany) according to the manufacturer’s protocol. Reverse-transcription polymerase chain reaction (RT-PCR) was performed using the PrimeScript RT Reagent Kit (Takara Bio Inc., Shiga, Japan) and Taq DNA polymerase (New England Biolabs, Ipswich, MA, USA). To quantify mRNA expression, real-time RT-PCR was performed using the StepOne™ Real-Time PCR System (Thermo Fisher Scientific Inc). All mRNA expression levels were normalized to those of 36B4 mRNA. The primer sequences are listed in Supplemental Table1.

### Assessment of cell viability

After pharmacological treatment, the cell viability was confirmed by Annexin V-FITC Apoptosis Detection Kit (Nacalai Tesque, Inc.). In detail, the cells were treated with trypsin-EDTA (Thermo Fisher Scientific Inc.), and the collected cells were washed with PBS. The cells (1 × 10^6^/100 μL) were incubated with Annexin V-FITC and Propidium iodide for 15 min at room temperature. Then, the cells were plated on glass-bottom dishes, and images were obtained using a microscope (ECLIPSE Ti2; Nikon Corporation) coupled with NIS elements v4.50 software.

### Statistical analysis

Data are presented as means ± S.E.M.. Student’s *t*-test and One-Way ANOVA analysis were used to identify statistically significant differences. Differences were considered significant at *p* < 0.05.

## RESULTS

### Methionine oxidant treatment enhanced the TRPV2 activity

First, we confirmed whether treatment with ChT (a methionine oxidant) at warm temperature activated mouse TRPV2 using Ca^2+^-imaging. Slight increases in [Ca^2+^]_i_ by 100 μM ChT treatment were observed in mock-transfected HEK293T cells at 32°C (Fig. 1A). Whereas significantly higher increases in [Ca^2+^]_i_ by 100 μM ChT treatment were observed in mTRPV2-expressing HEK293T cells at 32°C (Fig. 1B and E). This increase in [Ca^2+^]_i_ was significantly impaired when 10 μM SKF96365 (a TRPV2 inhibitor) or 10 μM ruthenium red (RR, a non-selective TRP channel inhibitor) was applied both before and during the treatment of 100 μM ChT (Fig. 1C-E). Unfortunately, while we checked the effect of 2-APB (a TRPV2 agonist) to confirm the functional expression of TRPV2, further increases in [Ca^2+^]_i_ by 500 μM 2APB treatment were rarely observed (Fig. 1C and D). Treatment with TRPV2 inhibitors for approximately 300 sec could affect TRPV2 activity before applying 2APB.

**Fig. 1.**
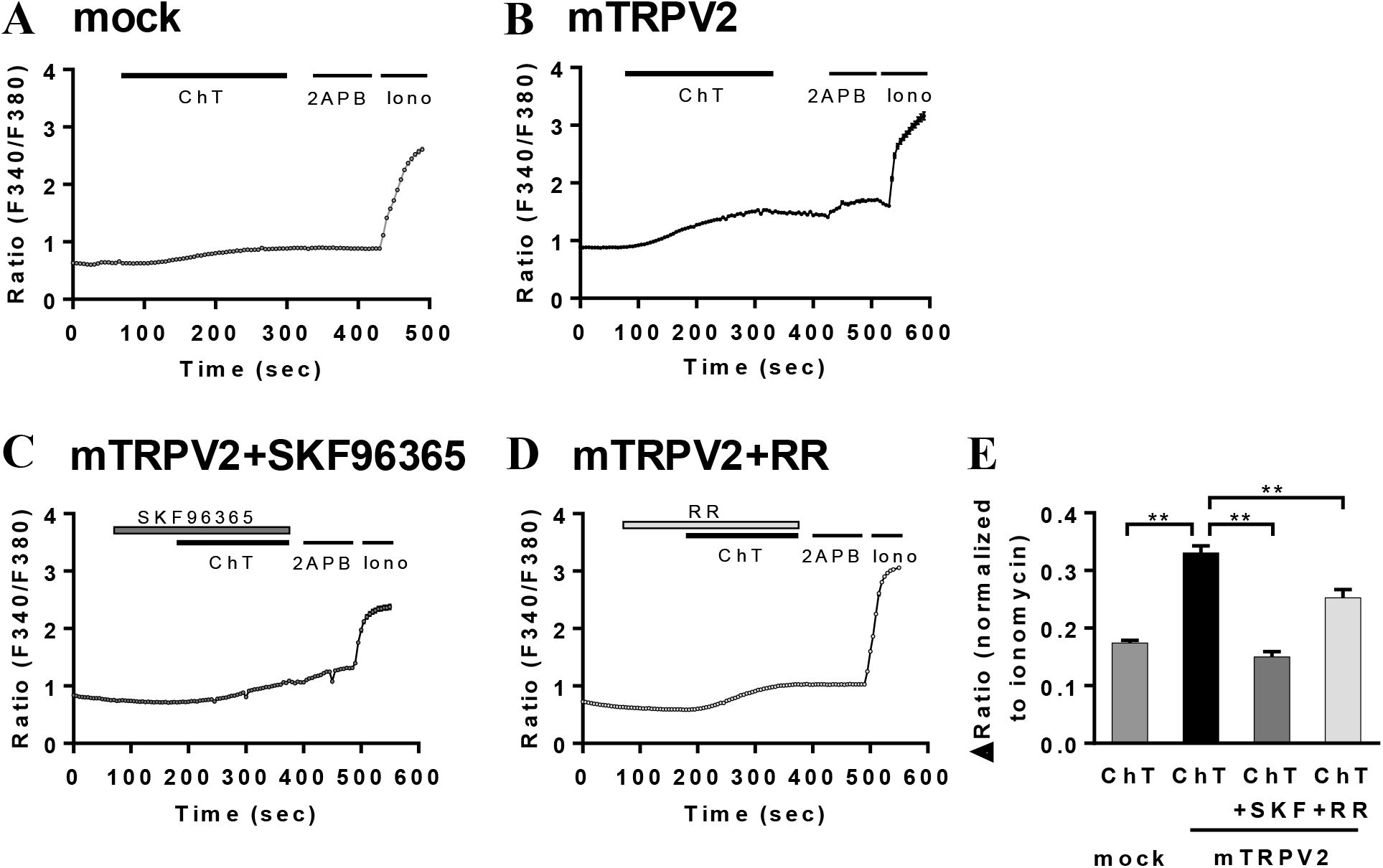
TRPV2 was activated by treatment of methionine oxidant in mTRPV2-HEK293T cells at 32°C. (A, B) Time course of intracellular Ca^2+^ concentration ([Ca^2+^]_i_) changes in response to 100 μM Chloramine-T (ChT) in mock-transfected HEK293T cells (A) and mTRPV2-expressing HEK293T cells (B). (C, D) Time course of [Ca^2+^]_i_ changes in response to 100 μM Chloramine-T (ChT) with 10 μM SKF96365 (a TRPV2 inhibitor; SKF, C) or 10 μM Ruthenium Red (a non-selective TRP channel inhibitor; RR, D) in mTRPV2-transfected HEK293T cells. (E) Summary of [Ca^2+^]_i_ increases by treatment of 100 μM ChT in mock-transfected HEK293T cells (mock) and mTRPV2-transfected HEK293T cells (mTRPV2) with or without inhibitors in mTRPV2-transfected HEK293T cells. 500 μM 2-Aminoethoxydiphenyl borate (2APB) was used as a positive control for TRPV2 and 5 μM Ionomycin (Iono) was used to confirm cell viability. Data are presented as mean ± SEM. n= 146-256. **;*p* < 0.01.

To determine whether TRPV2 is expressed in the brown adipocyte cell line, we measured TRPV2 gene expression levels. As shown in Figures 2A and B, while TRPV2 mRNA was detected in both pre-adipocytes and differentiated adipocytes, its expression in differentiated brown adipocytes was higher than that in pre-adipocytes. Using this brown adipocyte cell line, we investigated the effects of ChT on [Ca^2+^]_i_ changes in the differentiated brown adipocytes. Application of 100 μM ChT also exhibited increases in [Ca^2+^]_i_ at 32°C in brown adipocytes (Fig. 2C). Additionally, this increase in [Ca^2+^]_i_ was significantly impaired by co-application with 10 μM SKF96365 or 10 μM RR (Fig. 2D-F). Unfortunately, when we checked the effect of 2APB to confirm the functional expression of TRPV2, the increases in [Ca^2+^]_i_ by 500 μM 2APB treatment were not observed (Fig. 2D and E), as observed in mTRPV2-expressing HEK293T cells. These results indicate that methionine oxidation by ChT could activate TRPV2 at 32°C in differentiated brown adipocytes.

**Fig. 2.**
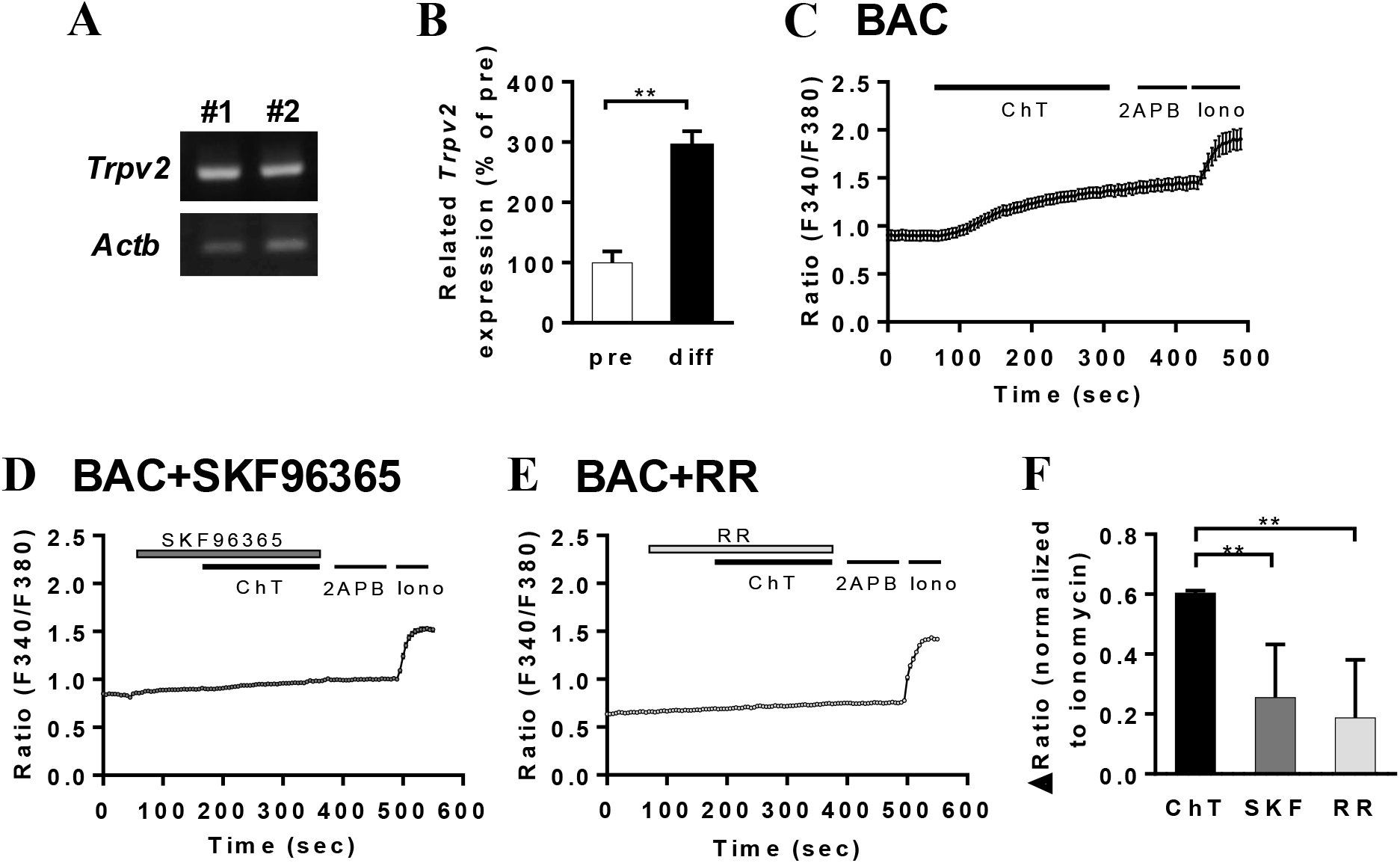
TRPV2 was activated by treatment of methionine oxidant in brown adipocytes at 32°C. (A) Confirmation of TRPV2 and *β*-actin mRNA expression in differentiated brown adipocytes by RT-PCR. (B) mRNA expression level of TRPV2 in pre-adipocytes (pre) and differentiated brown adipocytes (diff). mRNA expression level of TRPV2 was normalized to that of 36b4. Data are presented as mean ± SEM. n= 4-5. **;*P* < 0.01. (C) Time course of intracellular Ca^2+^ concentration ([Ca^2+^]_i_) changes in response to 100 μM Chloramine-T (ChT) in differentiated brown adipocytes. (D, E) Time course of [Ca^2+^]_i_ changes in response to 100 μM ChT with 10 μM SKF96365 (a TRPV2 inhibitor; SKF, D) or 10 μM Ruthenium Red (a non-selective TRP channel inhibitor; RR, E) in differentiated brown adipocytes. (F) Summary of [Ca^2+^]_i_ increases by treatment of 100 μM ChT with or without inhibitors in brown adipocytes. 1 mM 2-Aminoethoxydiphenyl borate (2APB) was used as positive control for TRPV2 and 5 μM Ionomycin (Iono) was used to confirm cell viability. Data are presented as mean ± SEM. n= 174-334. **;*p* < 0.01.

### TRPV2 activity was increased by methionine oxidant treatment in a temperature-dependent manner

We examined whether temperature changes can modulate the activity of TRPV2. We observed changes in [Ca^2+^]_i_ by temperature increase from 25°C to 38°C during application of 100 μM ChT in HEK293T expressing mTRPV2. As shown in Figures 3A and B, application of 100 μM ChT did not cause the increases in [Ca^2+^]_i_ at 25°C in both mock-transfected HEK293T cells and mTRPV2-expressing HEK293T cells. While elevating the temperature from 25°C to 38°C in the presence of 100 μM ChT elicited only minor increases in [Ca^2^_]_ in mock-transfected HEK293T cells (Fig. 3A), it induced much larger [Ca^2^_]_ rises in HEK293T cells expressing mTRPV2 (Fig. 3B, C). Furthermore, a temperature-dependent increase in [Ca^2+^]_i_ was also observed in the differentiated brown adipocytes (Fig. 4). These results suggest that TRPV2 can sense the physiological ranges of temperature changes by methionine oxidation.

**Fig. 3.**
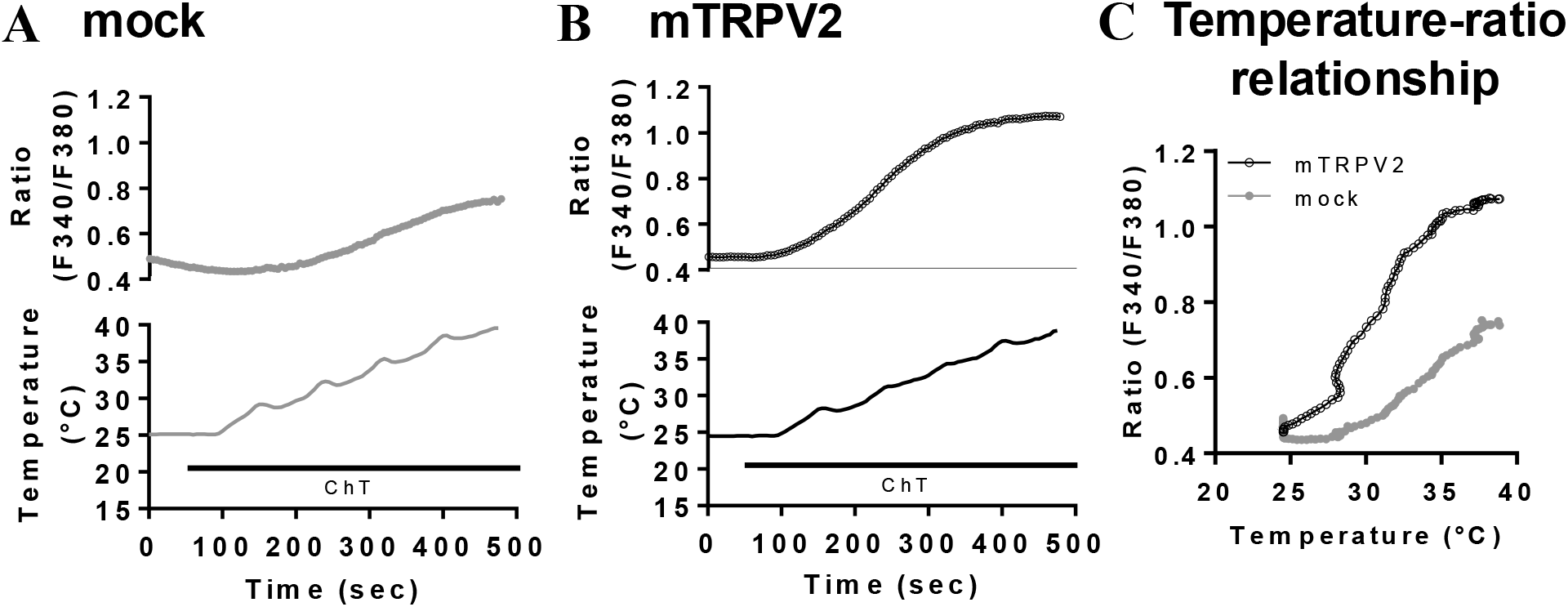
TRPV2 was activated by treatment of methionine oxidant in response to heat stimulation in mTRPV2-HEK293T cells. (A, B) Time course of intracellular Ca^2+^ concentration ([Ca^2+^]_i_) changes in response to 100 μM Chloramine-T (ChT) with temperature increase in mock-transfected HEK293T cells (A) and mTRPV2-transfected HEK293T cells (B). (C) Temperature–[Ca^2+^]_i_ profile from the traces in A and B. Data are presented as mean ± SEM. n= 178-256.

**Fig. 4.**
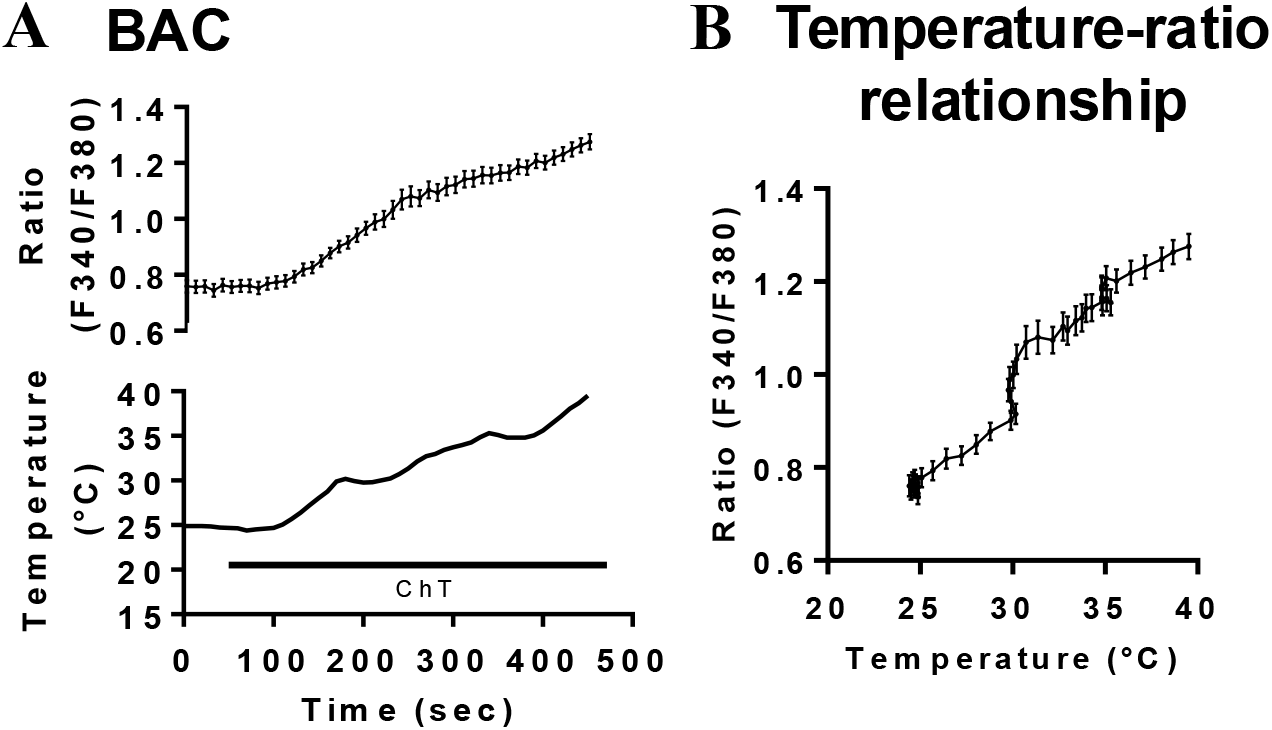
TRPV2 was activated by treatment of methionine oxidant in response to heat stimulation in brown adipocytes. (A) Time course of intracellular Ca^2+^ concentration ([Ca^2+^]_i_) changes in response to 100 μM Chloramine-T (ChT) with temperature increase in differentiated brown adipocytes. (B) Temperature– [Ca^2+^]_i_ profile from the traces in A. Data are presented as mean ± SEM. n= 43.

### Methionine oxidant treatment enhanced the expression of thermogenesis-related genes by β-adrenergic receptor agonist in brown adipocytes

Since we previously reported that TRPV2 positively regulates thermogenesis-related gene expression in brown adipocytes (Sun et al., 2016a), we examined whether the ChT treatment enhanced the expression of thermogenic genes in brown adipocytes. We measured the mRNA expression levels of thermogenesis-related genes stimulated by 3 μM Iso (a non-selective β-adrenergic receptor agonist) with or without 1 mM ChT for 4 hours in the differentiated brown adipocytes. As a result, the expression levels of *Ucp1* and *Pgc1a* in Iso- and ChT-treated brown adipocytes were significantly higher than those in Iso-treated brown adipocytes (Fig. 5A). In contrast, *Pparg* expression was not different between Iso-treated adipocytes and Iso- and ChT-treated adipocytes (Fig. 5A). We confirmed the cell viability against the treatment with Iso and ChT. The numbers of Annexin V-positive early apoptotic cells, Annexin V and PI-positive late apoptotic cells, and PI-positive dead cells were not significantly different between vehicle-treated adipocytes, and 3 μM Iso- and 1 mM ChT-treated adipocytes (Table 1), suggesting that Iso and ChT do not affect cell viability.

**Table 1.** Confirmation of the viability of Iso- and ChT-treated brown adipocytes.

| Treatment | Total cells | Annexin V | Propidium Iodide | Annexin V/<br>Propidium Iodide |
| --- | --- | --- | --- | --- |
| Control | 1611 | 61 (3.79 %) | 237 (14.71 %) | 4 (0.25 %) |
| 3 μM Iso + 1 mM ChT | 2454 | 94 (3.83 %) | 358 (14.59 %) | 7 (0.29 %) |

**Fig. 5.**
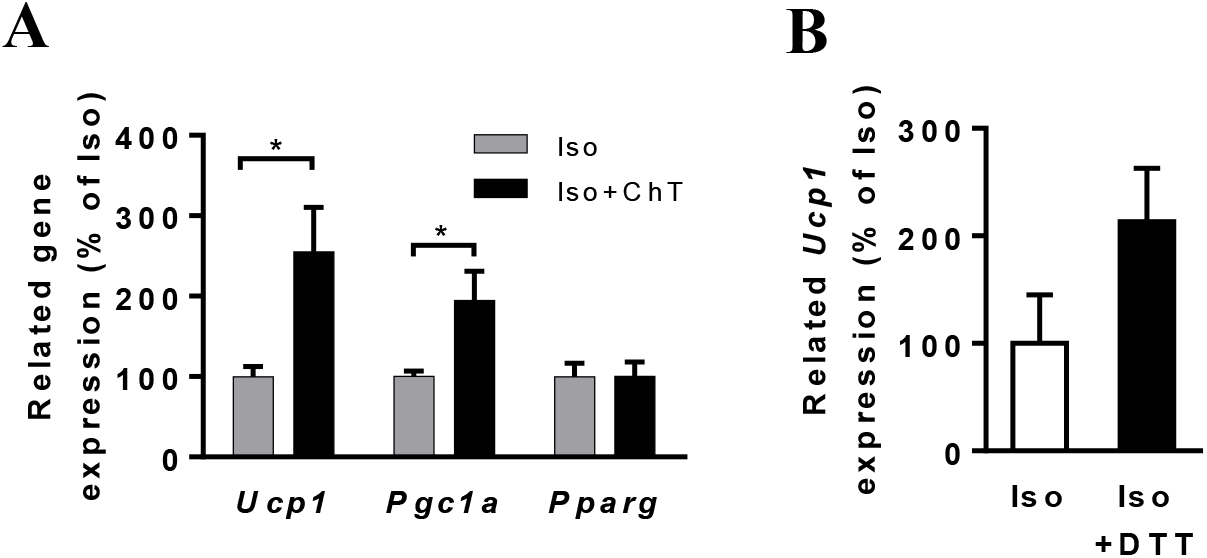
Treatment of methionine oxidant increased thermogenesis-related gene expression in brown adipocytes. (A) mRNA expression levels of UCP1, PGC1*α*, and PPAR*γ* in differentiated brown adipocytes in the presence of 3 μM Isoprenaline (Iso) with or without 1 mM Chloramine-T (ChT) for 4 hours. (B) UCP mRNA expression of differentiated brown adipocytes in the presence of 3 μM Iso treated with or without 3 mM dithiothreitol (DTT). mRNA expression levels of each gene were normalized to those of 36B4. Data are presented as mean + SEM. n= 5-7. *;*p*<0.05, **;*p*<0.01.

To check the involvement of endogenous ROS on thermogenesis-related gene expression through *β*-adrenergic receptor activation, we examined the effect of a reductant (Dithiothreitol, DTT) on Iso-induced increases in the expression of thermogenesis-related gene. Treatment with 3 mM DTT tended to enhance the expression of *Ucp1* in Iso-treated differentiated brown adipocytes (Fig. 5B). Therefore, we did not pursue the effect of DTT on ChT-treated enhancement of the expression of genes related to thermogenesis.

## DISCUSSION

Endogenously, TRPV2 is activated by mechanical stimuli and translocation of TRPV2 protein to the plasma membrane through PI3 kinase signaling (Aoyagi et al., 2010). Additionally, although TRPV2 is normally activated around high temperatures at 52°C, methionine oxidation of TRPV2 enhances its thermo-sensitivity, allowing TRPV2 to be activated at body temperature (Fricke et al., 2019). In this study, we found that treatment of ChT at 32°C caused [Ca^2+^]_i_ increase through TRPV2 activation in brown adipocytes (Fig. 2) and temperature elevation from 25°C to 38°C induced [Ca^2+^]_I_ increase in brown adipocytes treated with ChT (Fig. 4). These results suggest that TRPV2 might be activated by body temperature or BAT temperature by oxidation of the methionine residue of TRPV2. Moreover, our finding that treatment of ChT with *β*-adrenergic receptor agonist caused an increase in the expression of genes related to thermogenesis (Fig. 5A) may indicate the possibility that TRPV2 could positively regulate non-shivering thermogenesis by sensing BAT temperature. Our results also demonstrate that ChT treatment with Iso-treated condition did not affect the expression of *Pparg*, a marker for differentiation, in brown adipocytes (Fig. 5A). This result was consistent with our previous study, which showed no difference in *Pparg* expression in BAT between wild-type and TRPV2 knockout mice exposed to a cold environment (Sun et al., 2016a).

In this study, we found that methionine oxidation with *β*-adrenergic receptor activation enhanced the expression of the genes related to thermogenesis (Fig. 5A). Mitochondria constitute a significant source of ROS, generating superoxide and H_O_ from oxidoreductases associated with substrate catabolism and the electron transport chain (Brand, 2016). Some reports have indicated that ROS may activate thermogenesis by altering thermogenesis-related gene expression or by the direct redox modification of UCP1. Superoxide increases mitochondrial proton conductance by activating mitochondrial UCP1 (Echtay et al., 2002). Another report demonstrated that ROS were increased in the mitochondria by cold stimulation, contributing to the sulfenylation of UCP1, which induced the activation of UCP1-mediated thermogenesis (Chouchani et al., 2016). In addition, several TRP channels act as oxidative stress sensors (Ogawa et al., 2016; Jiang et al., 2023). Based on our results and previous reports, TRPV2 could be one of the targets of ROS produced during thermogenesis, and TRPV2 activation by ROS and tissue temperature positively regulates heat production in brown adipocytes. On the other hand, since it is also possible that *β*-adrenergic receptor activation could not be sufficient to oxidize TRPV2, we checked the effect of the antioxidant DTT on *β*-adrenergic receptor activation, as reported previously that DTT also reduced ChT-induced enhancement of thermo-sensitivity of TRPV2 (Fricke et al., 2019). Unfortunately, the application of DTT did not reduce *Ucp1* expression induced by the *β*-adrenergic receptor agonist. Instead, *Ucp1* expression tended to be increased upon co-application with DTT (Fig. 5B). Further analyses are necessary to understand the precise mechanisms of thermogenesis involving methionine oxidation.

We previously reported that an increase of [Ca^2+^]_i_ enhanced the expression of genes related to thermogenesis in brown adipocytes (Sun et al., 2016a). [Ca^2+^]_i_ increases through TRPV2 activation by methionine oxidation with warm temperatures may modulate gene expression. In addition, it was recently reported that [Ca^2+^]_i_ increases in brown adipocytes promote UCP1-independent and Ca^2+^-dependent non-shivering thermogenesis. Ca^2+^ transport contributes to non-shivering thermogenesis in BAT through sarco-endoplasmic reticulum ATPase activity (Ikeda and Yamada, 2020). It is possible that [Ca^2+^]_i_ increase through TRPV2 activation could also be involved in non-shivering thermogenesis via mechanisms other than an increase in thermogenesis-related gene expression.

In conclusion, methionine oxidation might activate TRPV2 at around body temperature and increase thermogenesis-related gene expression. Although it remains to be determined whether TRPV2 can be sufficiently activated by endogenous oxidative stress, it could be oxidized by mitochondria-derived ROS, leading to temperature sensing by TRPV2 in brown adipocytes.

## Conflict of interest statement

The authors declare that they have no conflicts of interest with the contents of this article.

## Funding and additional information

This research was supported by JSPS KAKENHI (Project No. JP23K05594 to K.U. and Project No. JP25K21079 to M.I.). This research was also supported in part by Takeda Science Foundation (K.U.), and a grant from Scientific Research on Innovative Areas ‘Thermal Biology’ (Project No. JP15H05928 to K.U.).

## Author contributions

M. I. and K. U. designed the experiments and wrote the manuscript. M. I. and K. U. performed the experiments and analyzed the data. M. I., S. K., T. G. and K. U discussed and interpreted the data. All authors read and approved the final manuscript.

## Data availability

All data are contained within the manuscript.

## Abbreviations

BAT: brown adipose tissue
cAMP: cyclic AMP
[Ca^2+^]_i_: intracellular Ca^2+^ concentration
ChT: Chloramine-T
diff: differentiated brown adipocytes
DTT: dithiothreitol
HEK293T: human embryonic kidney-derived 293T
H_2_O_2_: hydrogen peroxide
Iso: isoproterenol
mTPRV2: mouse transient receptor potential vanilloid 2
PGC1*α*: peroxisome proliferator-activated receptor gamma coactivator 1-alpha
PKA: protein kinase A
PPAR*γ*: peroxisome proliferator-activated receptor gamma
pre: pre-adipocytes
RT-PCR: reverse-transcription polymerase chain reaction
ROS: reactive oxygen species
RR: ruthenium red
SKF: SKF96365
TRPV2: transient receptor potential vanilloid 2
T_3_: triiodothyronine
UCP1: uncoupling protein 1
WAT: white adipose tissue
2APB: 2-aminoethoxydiphenylborane

## Notes

### Competing Interest Statement

The authors have declared no competing interest.

