## Supplemental Table 1 for "Methionine oxidation of TRPV2 regulates thermogenesis in brown adipocytes"

**Supplemental Table 1 Sequences of the primers for RT-PCR and RT-qPCR.**

| Gene | Purpose | Sequence |
| --- | --- | --- |
| *Trpv2* | RT-PCR/RT-qPCR | 5'- AGCACACAGGCATCTACAGTGTCA -3 |
|  |  | 5'- TTACTAGGGCTACAGCAAAGCCGA -3 |
| *Actb* | RT-PCR | 5'- TGTTACCAACTGGGACGACA -3 |
|  |  | 5'- AAGGAAGGCTGGAAAAGAGC -3 |
| *Ucp1* | RT-qPCR | 5'- TACCAAGCTGTGCGATGTCCA -3' |
|  |  | 5'- GCACACAAACATGATGACGTTCC -3' |
| *Pgc1a* | RT-qPCR | 5'- CCCTGCCATTGTTAAGACC -3' |
|  |  | 5'- TGCTGCTGTTCCTGTTTTC -3' |
| *Pparg* | RT-qPCR | 5'- GTGCCAGTTTCGATCCGTAGA -3 |
|  |  | 5'- GGCCAGCATCGTGTAGATGA -3 |
| *36b4* | RT-qPCR | 5'- GGCCCTGCACTCTCGCTTTC -3' |
|  |  | 5'- TGCCAGGACGCGCTTGT -3' |
